# Phantom auditory perception (tinnitus) is characterised by stronger anticipatory auditory predictions

**DOI:** 10.1101/869842

**Authors:** Marta Partyka, Gianpaolo Demarchi, Sebastian Roesch, Nina Suess, William Sedley, Winfried Schlee, Nathan Weisz

## Abstract

How phantom perceptions arise and the factors that make individuals prone to such experiences are not well understood. An attractive phenomenon to study these questions is tinnitus, a very common auditory phantom perception which is not explained by hyperactivity in the auditory pathway alone. Our framework posits that a predisposition to developing (chronic) tinnitus is dependent on individual traits relating to the formation and utilization of sensory predictions. Predictions of auditory stimulus frequency (remote from tinnitus frequency) were studied using a paradigm parametrically modulating regularity (i.e. predictability) of tone sequences and applying decoding techniques on magnetoencephalographic (MEG) data. For processes likely linked to short-term memory, individuals with tinnitus showed an enhanced anticipatory prediction pattern associated with increasing sequence regularity. In contrast, individuals without tinnitus engaged the same processes following the onset of the to-be-decoded sound. We posit that this tendency to optimally anticipate static and changing auditory inputs may determine which individuals faced with persistent auditory pathway hyperactivity factor it into auditory predictions, and thus perceive it as tinnitus. While our study constitutes a first step relating vulnerability to tinnitus with predictive processing, longitudinal studies are needed to confirm the predisposition model of tinnitus development.

## Introduction

Phantom perceptions do not require sensory input transduced by peripheral receptors. The common auditory phantom perception known as tinnitus affects approximately ~10% ^1,2^ of the population. Individuals experience tinnitus by consciously perceiving relatively simple sounds such as pure tones or narrow band noises without an identifiable objective environmental or bodily source. Tinnitus can be accompanied by substantial distress and reduced quality of life, which appears to be independent of the intensity of the perceived sound ^3^. The mechanisms by which this phantom sound emerges from ongoing brain activity (so-called “neural correlates”) have still not been resolved. A broad consensus supports the idea that some form of hearing damage (with or without clear audiometric changes) ^4–6^ stands at the outset of tinnitus development, leading to maladaptive functional or structural changes within or beyond the auditory system ^7–9^. By far the most popular view postulates a change of neural gain in deprived regions of the auditory pathway, thereby amplifying spontaneous activity which is interpreted as sound by downstream cortical regions (for review see ^10^; we will subsequently refer to this general idea as *altered gain model*).

Research along these lines has focused mostly on probable “neural correlate” candidates of tinnitus such as increased spontaneous firing rate or enhanced neural synchrony. The *altered gain model* of tinnitus is substantially supported by studies in animals ^11^, despite the obvious challenges in obtaining subjective reports. In humans the supporting evidence for this model is less apparent, partly because (contrary to animal models) the research is focused on chronic rather than acute tinnitus, but also due to a lack of understanding as to how measures commonly obtained in humans (such as oscillatory power in M/EEG or BOLD in fMRI) can be translated to those used to support the *altered gain model*. Based on human and animal works in other domains ^12^, reduced ongoing alpha or increased gamma in auditory regions pertinent for phantom sounds (for other auditory phantom percepts see ^13,14^) may be relevant to perception of tinnitus. However, the empirical evidence is inconclusive ^15,16^. With the exception of technical or practical issues that may complicate a convincing confirmation of the altered gain model in humans, other observations speak in favour of its explanatory insufficiency ^17^: 1) Only a fraction of individuals who suffer a hearing impairment will experience tinnitus (~70% following sensorineural hearing loss; see ^18–20^). 2) The onset of tinnitus and the onset of the hearing loss often occur at different times. 3) Not all cases of acute tinnitus will become chronic. One possibility to overcome these explanatory gaps is to frame tinnitus perception within a Bayesian inference framework ^21^, which emphasizes the constructive nature of perception being guided by internal models ^22^. In order to establish and improve internal models, incoming sensory input is compared to predictions (so-called *priors*), which need to be cast in real-time in dynamic environments. In a recent predictive coding view, tinnitus is seen as a consequence of a default prediction of silence altering to one of sound when faced with (enhanced) spontaneous activity (“tinnitus precursor”) along the auditory pathway ^21^. While conceptually overcoming many inconsistencies related to the altered gain model ^17^, strong support for this view is lacking partially due to the non-trivial task of deriving robust and direct measures of tinnitus-supporting priors from ongoing brain activity. Recent work has found indirect evidence of altered priors in established tinnitus^23^, but the question of how and why such altered priors should even emerge in certain individuals remains open.

A recent line of reasoning holds that increased precision of priors could drive hallucinatory experiences ^24,25^. Indeed, interindividual variability in prior strength assessed in a visuo-auditory conditioning task predicts the experience of hallucinations in daily life ^26^. We postulate that the predisposition to developing tinnitus may be contingent on an individual’s – putatively relatively stable, “trait-like” – tendency to more strongly engage in predictive processing in the auditory modality. Ideally individualized measures of auditory predictive processing tendencies would be obtained *before* a potentially tinnitus-inducing event and then compared between individuals that do or do not develop (chronic) tinnitus. However, this is difficult to pursue in humans for ethical and practical reasons. In a first step to establish our tinnitus-predisposition framework, we focus on comparing individuals with chronic tinnitus and healthy controls. Using stimulus frequencies remote from those of tinnitus should reduce the chance of identifying consequences rather than causes of tinnitus.

Our hypothesis implies that when processing auditory input, individuals with tinnitus should engage predictions more strongly, that is, either more accurately or anticipatory, compared with individuals without tinnitus. Recently we established a powerful experimental approach ^27^ showing in normal hearing individuals that more regular pure tone sequences activate tonotopically specific auditory templates in an anticipatory manner (see ^28,29^ for similar findings in the visual modality). In line with our predisposition framework, with increasing statistical regularities of sound sequences, individuals with tinnitus exhibited stronger anticipatory representations of upcoming stimuli.

## Results

34 individuals with chronic tinnitus (16 females) took part in the experiment. For 25 individuals in the Tinnitus group, age-matched volunteers without tinnitus (17 females) were recruited for the purpose of group comparisons. Magnetoencephalography (MEG) was used to record neural activity while participants passively listened to sequences composed of pure tones at four different carrier frequencies. High temporal expectation was ensured by a strict rhythmic presentation at 3 Hz. While sound onsets were perfectly predictable, the probability of *which* carrier frequency would be presented (and thus could be predicted) was varied by parametrically modulating the regularity (i.e. predictability) of sound sequences across conditions (see Figure 1a and Methods for details). To investigate feature-specific predictive auditory processing also in absence of stimulation, sounds were omitted randomly in 10% of presentations. Tinnitus characteristics and tinnitus-related distress were assessed with online versions of standardised questionnaires (see Methods for details) shortly prior to the visit to the laboratory.

**Figure 1:**
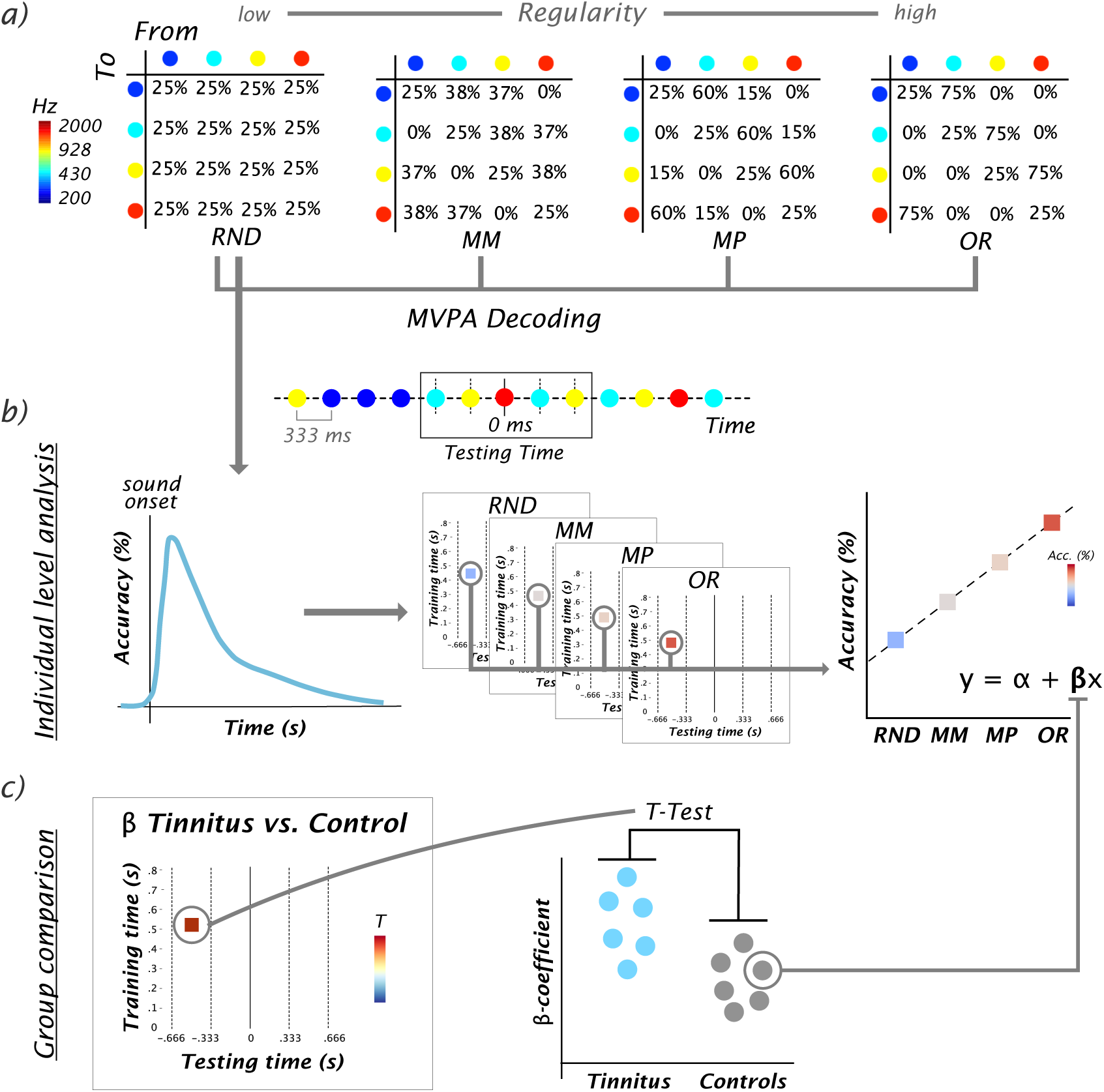
Experimental design and analysis rationale. a) Transition matrices used to generate sound sequences according to the different conditions (random [RD], midminus [MM], midplus [MP] and ordered [OR]) with a schematic example of a brief sound sequence. 10% of sound stimuli were randomly omitted. The “Testing Time” window corresponds to one trial with the to-be-decoded carrier frequency in the center (at 0 ms; marked by solid line), preceded and followed by two other tones (marked by dashed lines). b) For MVPA, time-shifted classifiers were trained on events in the random condition (left panel) and applied in a condition- and time-generalized manner to all conditions (middle panel). For every time-generalized data point, the dependence of decoding accuracy on the regularity of the sound sequence was quantified by a linear regression. c) At a group level, the resulting slopes (**β**-coefficients) of the regression analysis were compared between the tinnitus group and the control group.

To measure the dynamics of auditory predictions we used multivariate pattern analysis (MVPA) to derive feature (carrier frequency) specific information from the MEG data. Following our previous study ^27^, we trained classifiers to temporally decode the carrier frequency presented in the random sound sequence. These trained classifiers were subsequently tested on sound events in all regularity levels using time- and condition-generalization ^30^. For each individual we quantified how decoding accuracy was modulated by the regularity condition by extracting the slope (β coefficients) from a linear regression analysis. These were compared between the groups, yielding a time-generalized representation of T-values (see Figure 1).

### Normal neural encoding of carrier frequencies in tinnitus

Sensor level MEG data was used to decode the four carrier frequencies presented in the random sound sequence (Figure 1a and b). The trained classifiers were fundamental for targeting the main question of whether feature specific predictions in the auditory system are engaged differently in each of the groups in all further steps. In a first step, we could analyse the results of the simple decoding analysis for the random condition. Since this condition did not contain predictability-related information it allowed us to compare basic encoding of sound carrier frequencies in individuals with tinnitus with the control group. Both groups exhibited a rapid increase of decoding accuracy following sound onset robustly observed at an individual level (Figure 2a). Above chance (*p* < .05, Bonferroni corrected) decoding accuracy started immediately after stimulus onset in both samples (note that sampling rate was at 100 Hz). While peak increases were reached at approximately 100 ms, decoding accuracy remained statistically significant above chance for approximately ~500-600 ms with some interindividual variability. Remarkably, given the passive and non-engaging nature of the experiment, this means that carrier frequency specific information remained available during the two subsequent sound presentations. Interestingly, accuracy transiently increased approximately 100 ms after the subsequent stimulus onset (i.e. 450-500 ms after the to-be-decoded sound). Descriptively a similar pattern was observed following the next but one stimulus, albeit at a much smaller magnitude. These observations may reflect a sustained activation and reactivation of an auditory short-term memory trace enabling the formation of associations between events in temporal proximity, which is fundamental for subsequent learning of statistical regularities.

**Figure 2.**
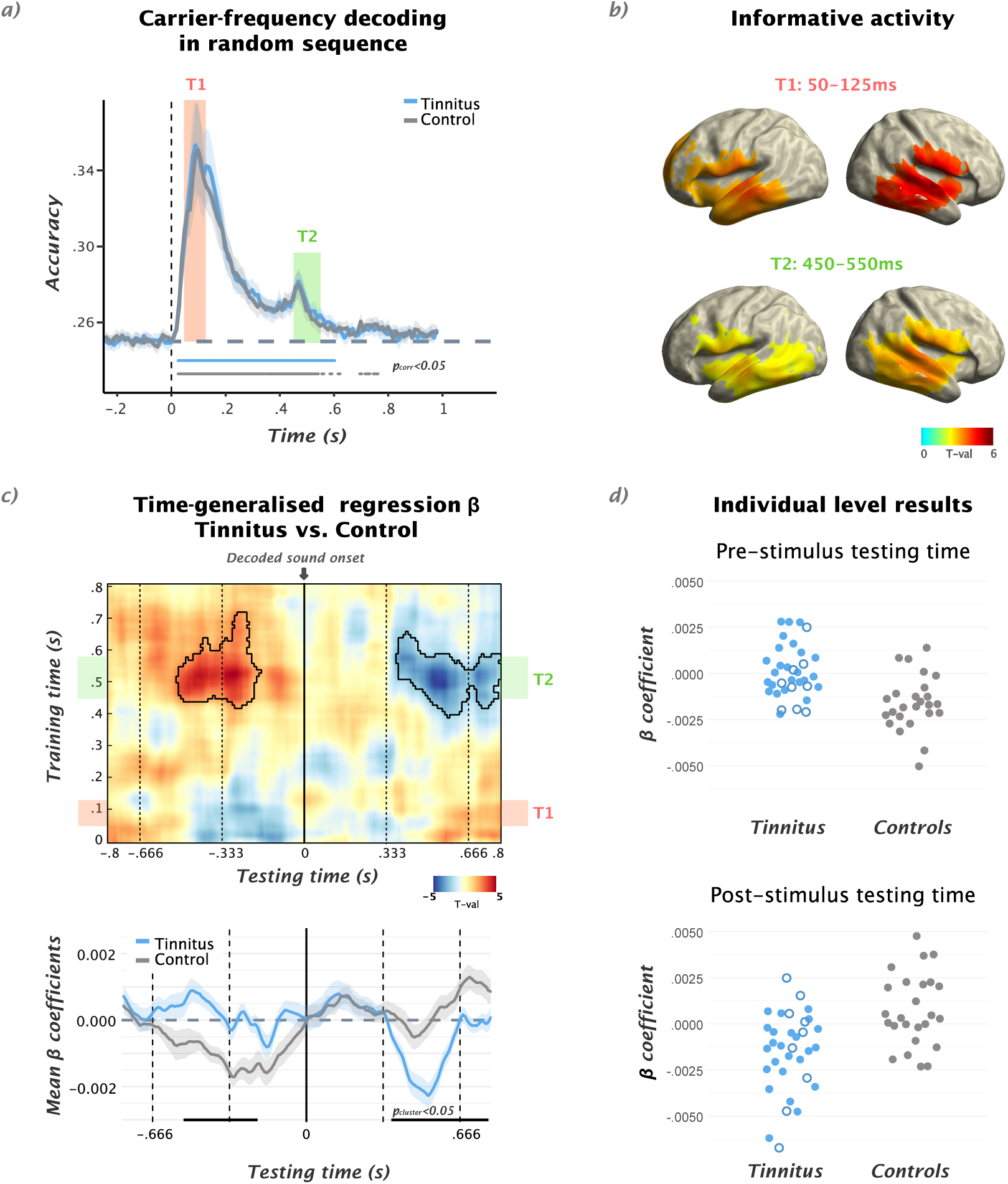
a) Temporal decoding of carrier frequencies in the random sound sequence for the tinnitus and control groups, respectively. In both groups, peak accuracy is reached after ~100 ms following sound onset. Above chance decoding accuracy is observed in a sustained manner up to ~600 ms (*p* < .05, Bonferoni corrected). No differences were observed between the groups. b) Source level depiction of Informative Activity for different periods: 50-125ms (T1) and 450-550ms (T2) after decoded sound presentation. The latter corresponds to the training time interval yielding pronounced group differences in the condition generalized analysis. c) (upper panel) Group comparison (see Figure 1c) of β-coefficient values between tinnitus vs. control groups in time-generalised matrix. Colors indicate t-values and solid black borders delimiting periods of significant difference (*p* < 0.05, cluster corrected). Lower panel: Time courses of β-coefficients averaged over 480-580ms training time-window, showing aforementioned effects driven by a relative increase of regularity-dependent carrier frequency specific activity prior to anticipated onset period and downregulation in the post-sound period in the tinnitus group. d) For illustration purposes, individual β-coefficient values within pre- and post-sound cluster are shown. While for the group comparison (shown in c) a subset of 25 individuals with tinnitus were taken into account, the full sample of 34 participants with tinnitus is displayed (individuals not considered in group comparison shown as hollow circles).

Importantly, we found no differences between the tinnitus group and the control group when carrier frequencies were presented randomly (Figure 2a). Since the upper carrier frequency of 2 kHz was at or below the audiometric edge for the majority of individuals with tinnitus (see audiograms in Supplementary Material Appendix 1 and 2), superior decoding results could plausibly be expected in the case of an enlarged neural representation of non-deprived tone frequencies resulting from tonotopic changes ^7^. Given the presence of hearing loss and potential tonotopic reorganization in individuals with tinnitus, the absence of a group difference in this simple carrier frequency decoding is of outstanding importance: that is, at a basic level individuals with tinnitus encode carrier frequencies equally well to individuals without tinnitus. This means that subsequently reported group differences are due to the manipulation of regularity (i.e. predictability) of the sound sequence.

### Regularity-driven carrier frequency specific neural information strongly differs between tinnitus and control groups

To adequately capture carrier-frequency specific, predictive-processing dynamics, we used a classifier trained on the random sound sequence (shown above) and applied it to all regularity levels in a time-generalized manner (Figure 1a). We used decoding accuracy as an indicator of the strength of internal representation of the particular stimulus frequency, and thus as a window into its utilisation in predictive processes. In order to quantify how the predictability of the carrier frequency modulates corresponding neural information, for each individual we calculated linear regressions (at each time-point over the entire temporal generalization matrix) between decoding accuracy and increasing regularity level (Figure 1b). In both groups, for the early training-time periods (~50-350 ms), similar patterns – in particular the anticipatory pre-activation of carrier frequency specific neural templates – were revealed as in the original experiment despite the slightly different analysis approach (see Supplementary Material Figure S3). For each point in the time-generalization matrix we compared the individual β-coefficients between groups using a t-test, reflecting differences in how carrier frequency specific predictions are modulated by the regularity of the sequence (Figure 1c).

Striking effects were obtained for relatively late training time intervals centred at around 530 ms. For trials in which the decoded sound was presented at testing time 0 (Figure 2c), we identified a positive cluster (*p* = 0.038) prior to the onset of the to-be-decoded event at approximately −530 ms to −200 ms, indicating a relatively stronger increase of decoding accuracy with regularity level for individuals with tinnitus. We interpret this as evidence of stronger correct anticipation of the present stimulus by individuals with tinnitus, in the higher regularity conditions where such anticipation is possible. We observed a similar effect in omission trials (see Supplementary Materials Figure S5). Time courses of β-coefficients averaged over the relevant late training time period (Figure 2c) showed that the intergroup differences were driven by opposing patterns: whereas individuals with tinnitus exhibited relatively increased carrier frequency specific information with stronger predictability prior to anticipated sound onsets, results for control individuals were marked by an augmenting absence (captured by the negative β-coefficients) of the carrier frequency pattern anticipated at 0 ms. Following the sound onset, a negative cluster (*p* = 0.05) between 360 ms and 800 ms was observed for the same training time interval. Similar to the prestimulus results, these post-sound onset effects are caused by inverse tendencies for the tinnitus and control groups (Figure 2c): that is, whereas individuals with tinnitus appeared to quickly deactivate carrier frequency patterns the more regular the sound sequence became, control individuals reactivate patterns of the decoded sound presented at 0 ms upon presentation of new events.

In order to make sense of this seemingly complex picture, it is important to detail the stimulation structure in light of our analysis approach, which focused on representation of the present stimulus frequency presented at time 0. Differing sequence regularities did not change the probability of the stimulus frequency remaining the same from one stimulus to the next (fixed at 0.25; i.e. diagonal of transition matrix), but increasing stimulus regularity did reduce the probability of the stimulus frequency remaining the same over separations of two or more stimuli. The observed regularity-related differences occurring from around two or more stimuli prior or subsequent to the present stimulus can be reconciled with the fact that relatively late training-time neural patterns capture this group-level effect. These patterns likely reflected processes associating sequential inputs, that is, short-term memory processes that integrate information over longer timescales. Our results suggest that in highly predictable sequences, control individuals engage these feature-specific auditory short-term processes in a more *reactive* way. Qualitatively this is similar to the manner they are activated in random sequences, that is, the stimulus that has just been heard is continuously represented and reactivated when new input arrives. Tinnitus individuals on the other hand exhibit a rather *proactive* engagement of the same processes with increasing regularity, preactivating stimulus representations in auditory short-term memory before their actual onset. Upon presentation of subsequent stimuli – which become less likely to be the same carrier frequency as presented at 0 – feature-specific neural patterns are downregulated. Overall, our results point to a dramatically altered involvement of higher level auditory short-term memory processes related to associating discrete events to form representations (“internal models”) of the statistical regularity of the sound sequence. These findings support the hypothesis that individuals with tinnitus utilize internal models in a more anticipatory manner when processing auditory events.

### Regularity-dependent engagement of internal models is unrelated to the magnitude of hearing loss and subjective tinnitus features

Following the demonstration of a marked group difference in activating late carrier frequency-specific neural patterns as a function of sequence regularity, we tested whether the magnitude of this process was related to subjectively rated tinnitus characteristics as well as audiometric features. Across the full (*N* = 34) tinnitus sample, we performed Spearman correlation between the averaged β-regression values corresponding in time to statistically significant anticipatory positive and post-stimulus negative clusters and magnitude of hearing loss (HLS, measured by Tinnitus Questionnaire), tinnitus loudness (TL) and tinnitus distress (TD) (see Supplementary Material Figure S5).

In spite of the explorative (liberal) testing without multiple comparison corrections, no significant correlation effects for any of these factors were identified for the prestimulus positive cluster (HLS: *rho* = −0.6, *p* = 0.75; TL: *rho* = −0.06, *p* = 0.73; TD: *rho* = 0.11, *p* = 0.53) nor for the post-stimulus negative effect (HLS: *rho* = −0.13, *p* = 0.45; TL: *rho* = −0.01, *p* = 0.95; TD: *rho* = −0.14, *p* = 0.43). The lack of relationships between prediction related neural effects with hearing loss add further support to the claim that the effects visible in group analysis are strictly regularity-dependent and not driven by low-level auditory processing. From a “neural correlate” perspective, the lack of correlation with tinnitus-specific (distress and loudness) measures would seem counterintuitive. However, this result is fully compatible with the predisposition view that we are advancing, proposing that individual predictive processing tendencies are relevant for the emergence and stabilization of tinnitus.

## Discussion

Current “neural correlate”-based approaches of tinnitus are insufficient to explain the interindividual varying trajectories that lead some individuals to develop (chronic) tinnitus following hearing damage but not others. Our predictive processing predisposition framework relies on inter-individual trait differences in applying internal models in the auditory system. Vulnerability to developing (chronic) tinnitus may arise from stronger tendencies to process incoming sounds according to internal model-based predictions: These tendencies could both refer to absolute strength (precision) or altered temporal dynamics (i.e. becoming more anticipatory) of auditory predictions. The individual’s predictive processing tendency could lead to different clinical outcomes when faced with potentially tinnitus-inducing events such as increased spontaneous activity and/or synchrony in the auditory pathway that follows hearing damage or noise overexposure. For instance, individuals better able to predict the dynamics of this spontaneous activity over time would form stronger predictions of it, thus facilitating its perception as an auditory entity through altered predictions^21^. However, other frameworks that emphasize the importance of top-down control of auditory activity to play a role in tinnitus generation (e.g. ^31^) are also compatible with our predisposition concept. In a first necessary step towards establishing support for this novel framework, we compared individuals with chronic tinnitus and controls without tinnitus, utilizing an approach ^27^ that allows us to scrutinize feature-specificity of predictive processes in the auditory system at high temporal resolution. In contrast to “neural correlate” approaches, no special importance was placed on the tinnitus frequency. Our main findings are: 1) basic processing of carrier frequencies is not altered in tinnitus; 2) higher-level (short-term memory-based) processing of carrier frequency exhibit a stronger anticipatory pattern in individuals with tinnitus as compared to controls; 3) the latter pattern is not correlated to factors such as magnitude of hearing loss or tinnitus-related variables (distress and loudness), in line with the idea that they reflect a more general predictive processing tendency of the individual.

Our approach to identifying modulation of feature-specific auditory activity as a function of predictability (set by the regularity of the sequence) used training classifiers to decode carrier frequencies in the random sound sequence. While our framework would predict strongest differences in situations when reliable internal models can be formed, it was important to also scrutinize processing of carrier frequencies when precise predictions cannot be made. Differences could be plausibly expected since most individuals with tinnitus exhibit some hearing loss at higher frequencies putatively leading to cortical reorganization: In particular an expanded representation of non-damaged cochlear regions ^7^ and potential improved sensory processing thereof ^32^ could imply an improved decoding performance in the random sequence. However, the temporal decoding patterns were virtually identical for both groups, with the characteristic features elaborated on in our previous report ^27^ (e.g. the rapid onset and relatively sustained above-chance decoding performance outlasting subsequent tone presentations). The lack of a group difference is overall in line with findings indicating no abnormal tonotopic representation in tinnitus ^33^ in contrast to earlier reports ^34^. Making a stronger point on this issue would require establishing that decoding performance in the random sequence can be taken as a quantitative proxy for tonotopic representation. Importantly for the current study, all group differences we reported result not from low-level, feedforward activation of tonotopically neural ensembles, but from adding varying levels of regularity to the sound sequence.

Indeed, striking regularity-dependent group differences were observed, with rich temporal information that can only be uncovered using high-temporal resolution methods: Firstly, while the general peak of decoding accuracy occurred at ~100 ms and in these early training-time windows exhibited a positive relationship with regularity (see Figure S3 in Supplementary materials) as described in ^27^, these early periods did *not* capture group differences. For late training-time intervals, however, marked group differences were observed. Interestingly, the relevant training time interval is ~150 ms after the onset of the sound following the to-be-decoded sound. This late increased accuracy for decoding carrier frequencies in the random sequence indicates a reactivation of a short-term memory representation of carrier frequency specific information presented at 0 ms (see also the descriptive similarity of Informative Activity patterns for early and late periods in Figure 2b). This process leads to a co-activation of new with previous input, which is crucial for associating discrete events via Hebbian principles. These learned associations are crucial for building up an internal model of the statistical regularities underlying the generation of the sound sequence. The selective involvement of these late processes in terms of group differences points to the role of high-level (memory based) auditory processes contributing to (or predisposing) tinnitus beyond purely bottom-up driven processes. An open question however remains as to whether these differences would be seen without the reactivation caused by a subsequent sound. A study systematically varying the ISI would be needed to resolve this issue, showing whether the latency of effects would remain relatively stable or follow the temporal separation of events.

Secondly, the temporal resolution of MEG allowed us to precisely describe the temporal dynamics of how these higher level auditory processes are engaged in the context of different levels of regularity of the sound sequence and how they differ between the groups (Figure 2c). Effects were dependent on whether the time-window of investigation (testing time) was prior to or following the onset of the to-be-decoded sound (testing time at 0 ms). Both groups showed an immediate engagement of these short-term memory-related auditory processes following the (perfectly predictable in time) sound onset. At later intervals (~600 ms), coinciding with the onset of the second sound following the to-be-decoded sound, decoding accuracy increased in the control group with increasing regularity of the sequence (see Figure 2c). The pattern was *reactive* in the sense that in periods prior to the anticipated onset of the to-be-decoded sound, carrier frequency specific information was less present with increasingly regular sounds. This indicates that short-term memory related auditory processes are engaged only once a predicted sound is presented, potentially contributing to a continuous update and stabilization of a formed internal model. Individuals with tinnitus, however, show an almost mirror-image pattern to the control group, with stronger anticipatory engagement of short-term memory related auditory processes when the sequence becomes more regular. Following the anticipated onset of a more predictable sound (carrier frequency) a marked disengagement of the relevant carrier frequency specific neural patterns is observed: this could be partially driven by processing the sound presented at 333 ms or anticipating the sound presented at 666 ms, both (usually) differing from the one presented at 0 ms in regular sequences. Irregardless, the results point to a dramatic difference with respect to internal models utilization between individuals with tinnitus and the control group. Overall, the more anticipatory pattern in tinnitus is in line with our belief that stronger predictive processing tendencies could identify individuals vulnerable to developing tinnitus. On a broader level the observed effects are also in accord with reports linking strong priors to general proneness to auditory hallucinations, even though a link between our data and those derived from computational modeling of behavioral data would need to be established. Also in contrast to a previous study supporting this notion ^26^, we derive our conclusions from neural data obtained during passive sound processing without experimentally inducing illusory percepts. The simplicity of our approach may be useful for studying altered predictive processing in other clinical groups, including ones in which behavioral assessment is challenging ^35^.

Albeit striking in terms of strength, the group effects reported here do not conclusively confirm a core idea that we are advancing, namely that increased internal model utilization tendencies in the auditory system predispose development of tinnitus. The absence of correlations with variables associated with tinnitus-induction (e.g. hearing loss) or consequences of tinnitus (e.g. loudness or distress), supports the view that the predictive processes we observe using our approach could be a temporally more stable “trait-like” feature of the individual. However, strong evidence would ultimately require longitudinal studies in humans ideally starting measurements prior to onset of (chronic) tinnitus, which is challenging (for an approach to inducing transient tinnitus see ^4^). Thus a next step may be to apply this paradigm in animal models of (chronic) tinnitus, where inter-animal variability has also been reported (e.g. ^36^). Such an approach should be relatively straightforward since the paradigm does not require any task for which the animal needs to be trained. Also when neural recording is performed using multiple electrodes, large parts of the analysis described here could be applied.

To summarize, we show for the first time enhanced anticipatory engagement of feature-specific high-level (putatively short-term memory based) predictive auditory processing in individuals experiencing chronically auditory phantom perception – tinnitus. However, whether this pattern constitutes a predisposing factor or is a consequence of tinnitus onset (despite being uncorrelated to tinnitus-relevant features) remains to be addressed in future studies. Resolving this issue has far-reaching consequences on a conceptual level by narrowing the explanatory gap of who will develop tinnitus following hearing damage. Also on a clinical level our work could have important implications, by potentially being able to identify individuals with greater risk of developing (chronic) tinnitus, thereby enabling more focused prevention or treatment efforts.

## Materials and methods

### Participants

A total of 34 individuals with tinnitus (17 females, 20-67 years old, *mean* age=45.12, *sd*=13.65) participated in the experiment: 25 (16 females, 20-66 years old, *mean* age= 40.92, *sd*=13.17) were age-matched (in all cases but one both age- and sex-matched) with the control group and used for group comparisons. Tinnitus related questionnaires (German version of *Tinnitus Questionnaire*, TQ; ^37^, *Tinnitus Sample Case History Questionnaire*, TSCHQ^41^ and 10-point scale *Tinnitus Severity*, TS) were collected for individuals with tinnitus. Standardized pure-tone audiometric testing for frequencies from 125Hz to 8kHz was performed in 31 out of 34 tinnitus participants using Interacoustic AS608 audiometer. 25 volunteers (17 females, 21-65 years old, *mean* age=41.56, *sd*=13.68) reporting no relevant audiological, neurological or psychiatric treatment history took part as a control group. 12 of the group were part of an experiment published elsewhere ^27^. Control subjects were age-matched to each tinnitus participant by the +/-3 years criterion, selecting the closest match in cases where more than one subject was eligible. No differences were shown for age between the samples comprised in the intergroup analysis (*t* = 0.17, *p* = 0.89). All participants provided written informed consent prior to participating. The experimental protocol was approved by the ethics committee of the University of Salzburg (EK-GZ: 22/2016 with Addenda).

### Stimuli and experimental procedure

Five head position indicator (HPI) coils were applied on the scalp of the subjects prior to entering the MEG shielded chamber. The Polhemus FASTRAK (Polhemus, Colchester, Vermont, U.S.A) digitizer was used to digitize head shape and position for each individual via marking of anatomical references (nasion and left/right pre-auricular points), location of HPI coils and approximately 300 additional points over the scalp. Before the start of the actual paradigm, a 5 min resting state recording was performed (not reported here), when subjects were asked to simply look at the center of the rear-projection screen.

During the experiment, participants watched a silent movie (“Cirque du Soleil: Worlds Away”), while passively being exposed to different tone sequences (Figure 1a). No instruction considering the sound stimuli was provided. The movie was displayed on the screen inside the shielded room using a projector (PROPIXX, VPixx technologies, Canada) and a periscope, whereas auditory stimulation was delivered to both ears via MEG-compatible pneumatic in-ear headphones (SOUNDPixx, ibid). Four different pure (sinusoidal) tones were presented, with carrier frequencies logarithmically spaced between 200 to 2000 Hz (200 Hz, 431 Hz, 928 Hz, 2000 Hz). Each of the tones lasted 100 ms, tapered with 5 ms linearly ascending/descending periods at both ends. Sounds were presented at a constant 3Hz stimulation rate.

Each participant was presented four blocks of tone sequences comprising 4000 stimuli, each lasting approximately 22 mins. The number of particular tone frequencies was balanced across blocks, so the condition-blocks varied solely by presentation order, which was parametrically modulated in their regularity (entropy) level using different transition matrices ^42^. In the random condition (RD, highest entropy or lowest regularity; see Figure 1a) there was an equal transition probability from one sound to another (thus preventing any possibility of accurately predicting an upcoming stimulus). Conversely, in the ordered condition (OR, lowest entropy level or highest regularity), presentation of one sound was for the majority (75% of cases) systematically followed by the particular other sound. Additionally, two intermediate entropy conditions were included, labelled here as midminus (MM) and midplus (MP). To control for the influence of self-repetitions, the diagonal of the transition matrices was set to be always 25% across all entropy conditions. The experiment was written using the MATLAB (ver. 9.1 The MathWorks, Natick, Massachusetts, U.S.A) based Psychophysics Toolbox ^43^.

### MEG data acquisition and preprocessing

Brain magnetic activity was measured using a whole-head MEG (Triux, MEGIN Oy, Finland), sampling the signal at 1000 Hz and with the default hardware filters set by the manufacturer (0.1 Hz high pass – 330 Hz low pass). Subjects were comfortably seated inside a dimly lit magnetically shielded room (AK3b, Vacuumschmelze, Germany). Signals were captured by 102 magnetometers and 204 planar gradiometers placed in 102 different positions. We used a signal space separation algorithm (SSS^44^) implemented in the Maxfilter program (version 2.2.15) to attenuate external noise from the MEG signal (mainly 16.6Hz, and 50Hz plus harmonics) and realign data to a common standard head position (“-trans default” Maxfilter parameter) across different blocks based on the measured head position at the beginning of each block ^45^. The rest of the subsequent analysis was performed on magnetometers only, given the mixing of information between the two sensors types after the Maxfilter step ^46^.

Data analysis was carried out with scripts written in-house, using the Fieldtrip toolbox ^47^ (git version 20170919). First a high-pass filter at 0.1 Hz (6th order zero-phase Butterworth filter) was applied to the raw data. Then, the continuous data were chunked in 10 s blocks, down-sampled to 256 Hz, and used as input to an Independent Component Analysis (ICA) algorithm. The ICA components were visually inspected to find eye blinks, eye movements, heartbeat and 16⅔ Hz (German/Austrian train power supply) artifacts. Finally, the continuous data were epoched from 1 s before to 1 s after target sound/omission onset and the artifactual components projected out (mean 3.6 ± 1.2 SD) components removed on average per each subject). All trials were kept using these preprocessing steps^45^. A further 30 Hz low pass filter (6th order zero-phase Butterworth filter) and 100 Hz resampling were applied to the epochs, before continuing with the multivariate pattern analysis (MVPA).

### Multivariate Pattern Analysis (MVPA) and classifier weights projection

We used MVPA as implemented in the MVPA-Light (https://github.com/treder/MVPA-Light, commit 003a7c), forked and modified in order to extract the classifier weights (https://github.com/gdemarchi/MVPA-Light/tree/devel). In essence, we implemented the analysis of carrier frequency decoding separately for sound and omission trials (*sound-to-sound* decoding and *sound-to-omission* decoding, respectively). We defined four targets (classes) for the decoding related to the carrier frequency of the sound presented in each trial. In order to focus solely on neural templates corresponding to carrier frequency-related information and avoid any potential carry over effect from the previous sound, the classifier was trained only on the random (RD) sounds and the preceding tone frequencies were balanced across trials. The exact details of the MVPA analysis have been described elsewhere ^25^. An identical procedure was applied to sound-to-omission decoding (see Figure S4 in Supplementary materials). We trained a multiclass LDA classifier on each sample point of the random (RD) condition and tested on all regularity level conditions for each time point of the testing set using a temporal generalization method ^30^. This enabled classifiers to generalize to each point in a time-shifted manner. Given the cross decoding nature of this approach, no cross-validation was performed, except for the testing on random (RD) tones, where a 5-fold cross validation, repeated five times, was implemented. For the sound-to-sound and sound-to-omission decoding, time generalization was calculated for each entropy level separately, resulting in four generalization matrices, one for each entropy level. For each subject, classification accuracy was then averaged at the group comparison level. Finally, and mainly for depiction purposes, the training decoders weights were extracted and projected in the source space, to localize the informative activity (see Figure 2b) related to carrier-frequency processing ^27,48^.

### Statistical analysis

As a first step, we extracted the dependence on entropy level within tinnitus and control groups. We arranged accuracy results for sounds from random to ordered and we then computed a regression for each single point of the testing-training 2D accuracy matrices, using the MATLAB built in least square *mldivide* algorithm (“\”), resulting in a training time by testing time matrix of slopes (“β”) for each subject, discarding intercepts. To compare the groups (25 Tinnitus subjects vs 25 age matched controls), we ran a t-test between the two matrices with coefficients obtained in the regression step, inputting them in the form of time-frequency 2D structures (time-generalised β values) in the ft_freqstatistics fieldtrip function. In order to account for multiple comparisons, we used a nonparametric cluster permutation test ^49^, with 1000 permutations and a *p* < 0.05 to threshold the clusters.

We pursued further analysis with questionnaire data using R ^50^. In the whole sample of participants with tinnitus (*Tinnitus Ws*, N=34) we performed a Spearman correlation of the β-coefficient values corresponding to the time-point of the maximum and the minimum t-value in intergroup analysis (comprised in positive and negative significant clusters emerging in group comparison for sound trials, see Figure 2c) with hearing loss (averaged audiogram for both ears), tinnitus loudness (10-point scale) and tinnitus distress scores (TQ). (see Supplementary Material Figure S5).

## Acknowledgements

We thank Mr. Manfred Seifter for the help with the measurements and Miss Hayley Prins for proofreading it.

This research is a part of the European School for Interdisciplinary Tinnitus Research project and has received funding from the European Union’s Horizon research and Innovation programme under the Marie Sklodowska-Curie grant agreement number 722046.

## Supplementary Materials

**Table S1.**
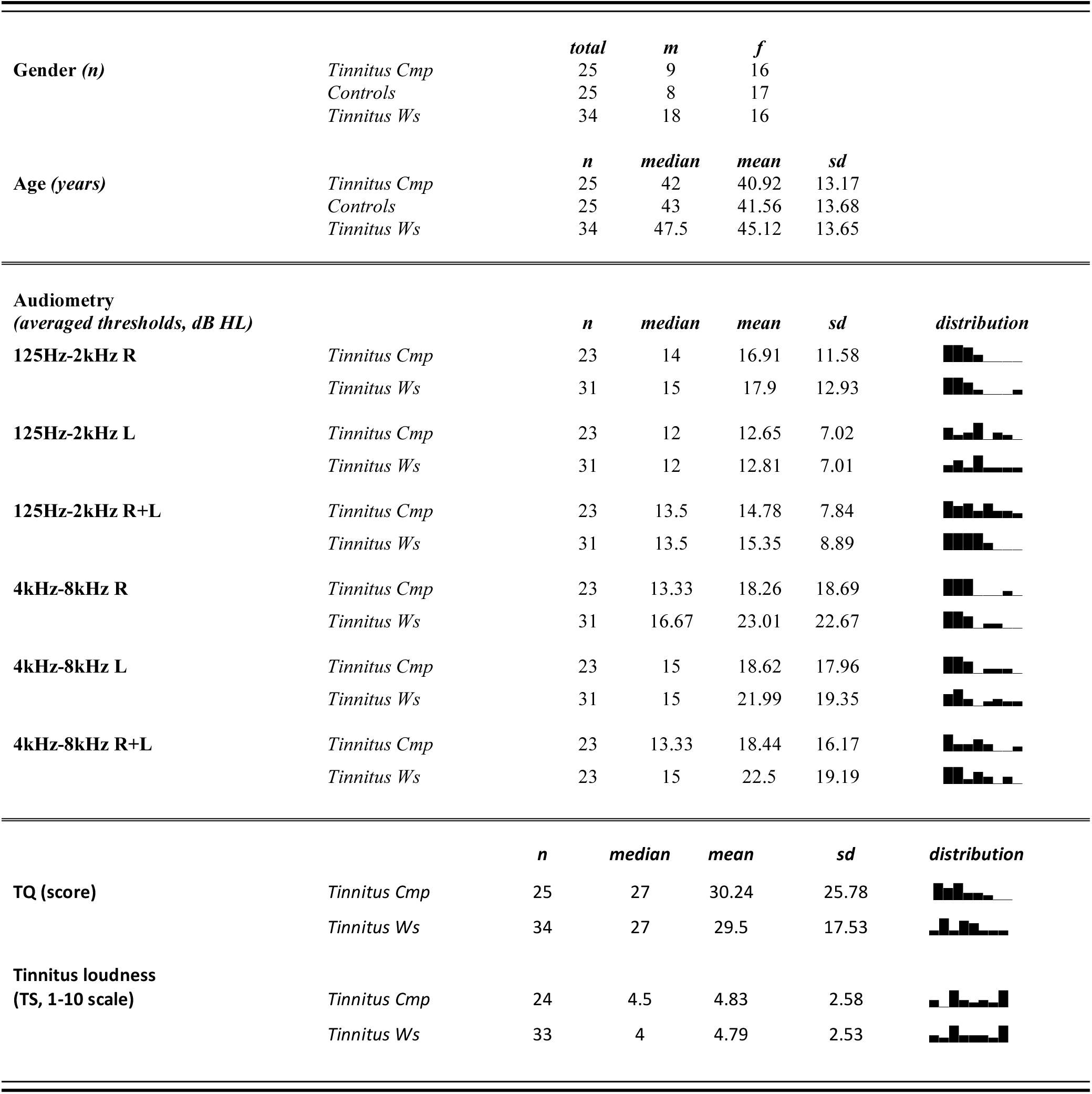
Demographic characteristics of the subject sample and descriptive statistics for averaged hearing loss and tinnitus characteristics in Tinnitus groups: *Tinnitus Cmp* for participants included in group comparison with Controls, *Tinnitus Ws* for the whole sample of subjects with tinnitus. Standard pure-tone audiogram values were averaged for each individual for the lower (125Hz-2kHz) and higher (4kHz-8kHz) frequency bands and presented here for right (R), left (L) and both ears (R+L). Tinnitus distress scores presented for Tinnitus Questionnaire (TQ). Tinnitus Loudness reported on the scale 1-10 from the Tinnitus Severity questionnaire.

**Table S2 a.** Detailed audiograms (in dB HL) for each subject with tinnitus, right ear.

| **Subject No** | **Group Comparison** | **125 Hz**<br>**R** | **250 Hz**<br>**R** | **500Hz**<br>**R** | **1kHz**<br>**R** | **2kHz**<br>**R** | **4kHz**<br>**R** | **6kHz**<br>**R** | **8kHz**<br>**R** |
| --- | --- | --- | --- | --- | --- | --- | --- | --- | --- |
| **1** | Yes | 10 | 15 | 10 | 5 | 0 | 5 | 5 | 15 |
| **2** | Yes | 5 | 5 | 5 | 5 | 5 | 0 | 0 | 5 |
| **3** | Yes | -5 | 15 | 10 | 5 | 10 | 10 | 20 | 10 |
| **4** | Yes | 5 | 5 | 5 | 5 | -5 | 0 | -5 | 10 |
| **5** | Yes | 15 | 20 | 20 | 5 | 10 | 10 | 10 | 10 |
| **6** | Yes | 15 | 15 | 10 | 10 | 5 | 5 | 10 | 5 |
| **7** | Yes | *na* | *na* | *na* | *na* | *na* | *na* | *na* | *na* |
| **8** | Yes | 5 | 5 | 15 | 10 | 10 | 5 | 5 | 5 |
| **9** | Yes | *na* | *na* | *na* | *na* | *na* | *na* | *na* | *na* |
| **10** | Yes | 25 | 25 | 25 | 15 | 15 | 10 | 20 | 20 |
| **11** | Yes | 35 | 50 | 50 | 45 | 15 | 0 | 0 | 0 |
| **12** | Yes | 25 | 15 | 15 | 10 | 10 | 5 | -10 | 0 |
| **13** | Yes | 15 | 10 | 10 | 10 | 20 | 25 | 30 | 15 |
| **14** | Yes | 10 | 10 | 10 | 10 | 0 | 15 | 20 | 30 |
| **15** | Yes | 20 | 25 | 20 | 20 | 20 | 15 | 20 | 10 |
| **16** | Yes | 20 | 25 | 25 | 35 | 30 | 30 | 25 | 10 |
| **17** | Yes | 10 | 0 | 10 | 10 | 20 | 15 | 20 | 0 |
| **18** | Yes | 45 | 65 | 55 | 55 | 50 | 55 | 75 | 80 |
| **19** | Yes | 20 | 15 | 20 | 15 | 10 | 20 | 40 | 25 |
| **20** | Yes | 30 | 25 | 15 | 10 | 10 | 0 | 5 | 5 |
| **21** | Yes | 5 | 5 | 5 | 30 | 15 | 25 | 45 | 45 |
| **22** | Yes | 25 | 25 | 20 | 5 | 0 | 10 | 20 | 15 |
| **23** | Yes | 10 | 15 | 15 | 10 | 5 | 10 | 35 | 5 |
| **24** | No | 20 | 20 | 20 | 20 | 20 | 55 | 60 | 45 |
| **25** | Yes | 25 | 25 | 25 | 30 | 20 | 40 | 60 | 70 |
| **26** | Yes | 15 | 15 | 25 | 15 | 10 | 15 | 35 | 15 |
| **27** | No | 10 | 5 | 5 | 10 | 10 | 55 | 60 | 55 |
| **28** | No | 25 | 25 | 25 | 20 | 20 | 20 | 15 | 20 |
| **29** | No | *na* | *na* | *na* | *na* | *na* | *na* | *na* | *na* |
| **30** | No | 30 | 30 | 20 | 20 | 30 | 40 | 55 | 60 |
| **31** | No | 10 | 10 | 10 | 0 | 5 | 5 | 5 | -5 |
| **32** | No | 25 | 25 | 20 | 20 | 25 | 20 | 25 | 30 |
| **33** | No | 15 | 10 | 15 | 5 | 10 | 10 | 15 | 10 |
| **34** | No | 50 | 55 | 60 | 60 | 70 | 80 | 90 | 100 |

**Table S2 b.** Detailed audiograms (in dB HL) for each subject with tinnitus, left ear.

| *Subject No* | *Group Comparison* | *125 Hz*<br>*L* | *250 Hz*<br>*L* | *500Hz*<br>*L* | *1kHz*<br>*L* | *2kHz*<br>*L* | *4kHz*<br>*L* | *6kHz*<br>*L* | *8kHz*<br>*L* |
| --- | --- | --- | --- | --- | --- | --- | --- | --- | --- |
| **1** | Yes | 0 | 5 | 5 | 0 | -5 | 0 | 0 | 0 |
| **2** | Yes | 5 | 5 | 10 | 0 | 0 | 0 | -5 | 0 |
| **3** | Yes | 5 | 10 | 10 | 10 | 0 | 10 | 10 | 15 |
| **4** | Yes | 5 | 5 | 5 | 5 | 0 | 0 | -5 | -5 |
| **5** | Yes | 25 | 25 | 20 | 15 | 10 | 10 | 15 | 10 |
| **6** | Yes | 10 | 15 | 10 | 5 | 0 | 5 | 0 | 5 |
| **7** | Yes | *na* | *na* | *na* | *na* | *na* | *na* | *na* | *na* |
| **8** | Yes | 0 | 5 | 10 | 0 | 5 | 5 | 5 | 5 |
| **9** | Yes | *na* | *na* | *na* | *na* | *na* | *na* | *na* | *na* |
| **10** | Yes | 30 | 30 | 25 | 15 | 15 | 20 | 55 | 75 |
| **11** | Yes | 40 | 10 | 10 | 5 | 5 | 10 | 15 | 25 |
| **12** | Yes | 25 | 20 | 15 | 0 | 0 | 0 | -5 | 0 |
| **13** | Yes | 15 | 10 | 15 | 10 | 10 | 5 | 25 | 20 |
| **14** | Yes | 15 | 15 | 15 | 10 | 0 | 15 | 20 | 25 |
| **15** | Yes | 20 | 20 | 20 | 20 | 20 | 10 | 15 | 20 |
| **16** | Yes | 20 | 25 | 25 | 30 | 40 | 50 | 35 | 40 |
| **17** | Yes | 5 | 5 | 10 | 15 | 25 | 10 | 20 | 15 |
| **18** | Yes | 5 | 5 | 15 | 10 | 10 | 5 | 15 | 10 |
| **19** | Yes | 20 | 15 | 25 | 25 | 15 | 30 | 35 | 25 |
| **20** | Yes | 15 | 15 | 15 | 10 | 5 | 0 | 5 | 10 |
| **21** | Yes | 10 | 5 | 5 | 0 | 10 | 25 | 35 | 55 |
| **22** | Yes | 20 | 15 | 15 | 10 | 0 | 5 | 15 | 10 |
| **23** | Yes | 5 | 5 | 5 | 10 | 10 | 10 | 5 | 15 |
| **24** | No | 15 | 20 | 20 | 25 | 35 | 60 | 55 | 60 |
| **25** | Yes | 20 | 10 | 10 | 10 | 10 | 20 | 40 | 80 |
| **26** | Yes | 5 | 10 | 15 | 15 | 15 | 20 | 45 | 55 |
| **27** | No | 10 | 10 | 5 | 5 | 5 | 40 | 60 | 55 |
| **28** | No | 20 | 20 | 20 | 10 | 5 | 5 | 5 | 10 |
| **29** | No | *na* | *na* | *na* | *na* | *na* | *na* | *na* | *na* |
| **30** | No | 25 | 20 | 15 | 15 | 35 | 50 | 60 | 60 |
| **31** | No | 5 | 5 | 10 | 10 | 0 | 5 | 5 | 5 |
| **32** | No | 25 | 25 | 20 | 10 | 5 | 20 | 30 | 35 |
| **33** | No | 10 | 15 | 10 | 10 | 15 | 30 | 35 | 55 |
| **34** | No | 30 | 30 | 30 | 20 | 20 | 35 | 45 | 55 |

**Figure S3.**
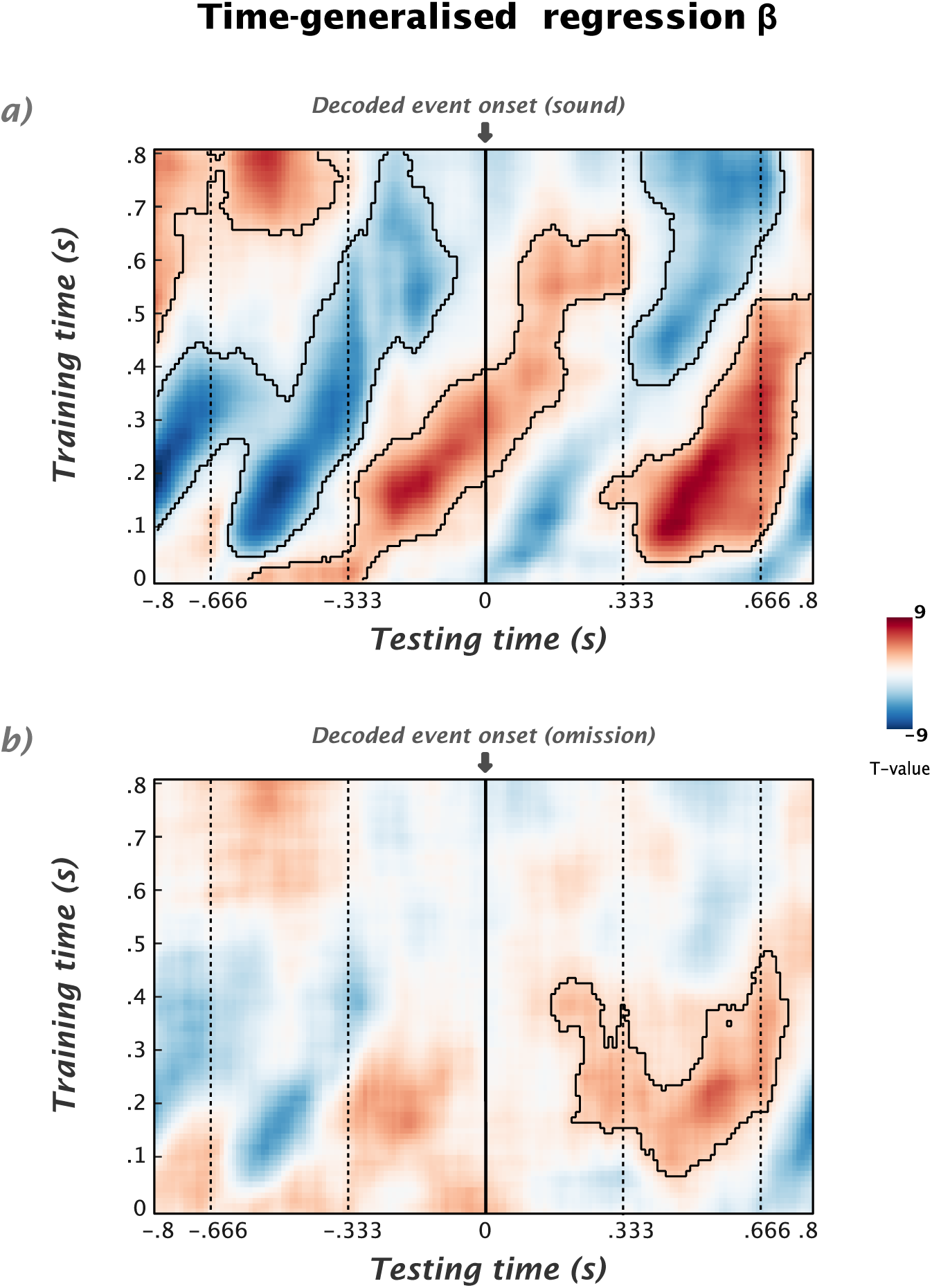
Time generalisation of β-coefficient values in the sample of Tinnitus Cmp and Controls joint together (N=50), tested against 0: a) For sound trials, we observe a pattern as reported in the previous study: carrier frequency specific templates are most strongly driven by early training time (~100-150 ms) and emerge in accordance to regularity level, in anticipatory period before the presentation of the sound as well as after the presentation of the consecutive tone (~450 ms in testing time). In the omission trials (b) the pre-stimulus effects do not reach significance but we observe the post-stimulus significant linear increase of decoding accuracy with regularity, emerging at approx ~150 ms after expected sound omission.These results point to presence of anticipatory activation of the templates corresponding to carrier frequency dependent on predictability and reactivation based on the knowledge about the sound sequence, putatively related to short-term memory processes.

**Figure S4.**
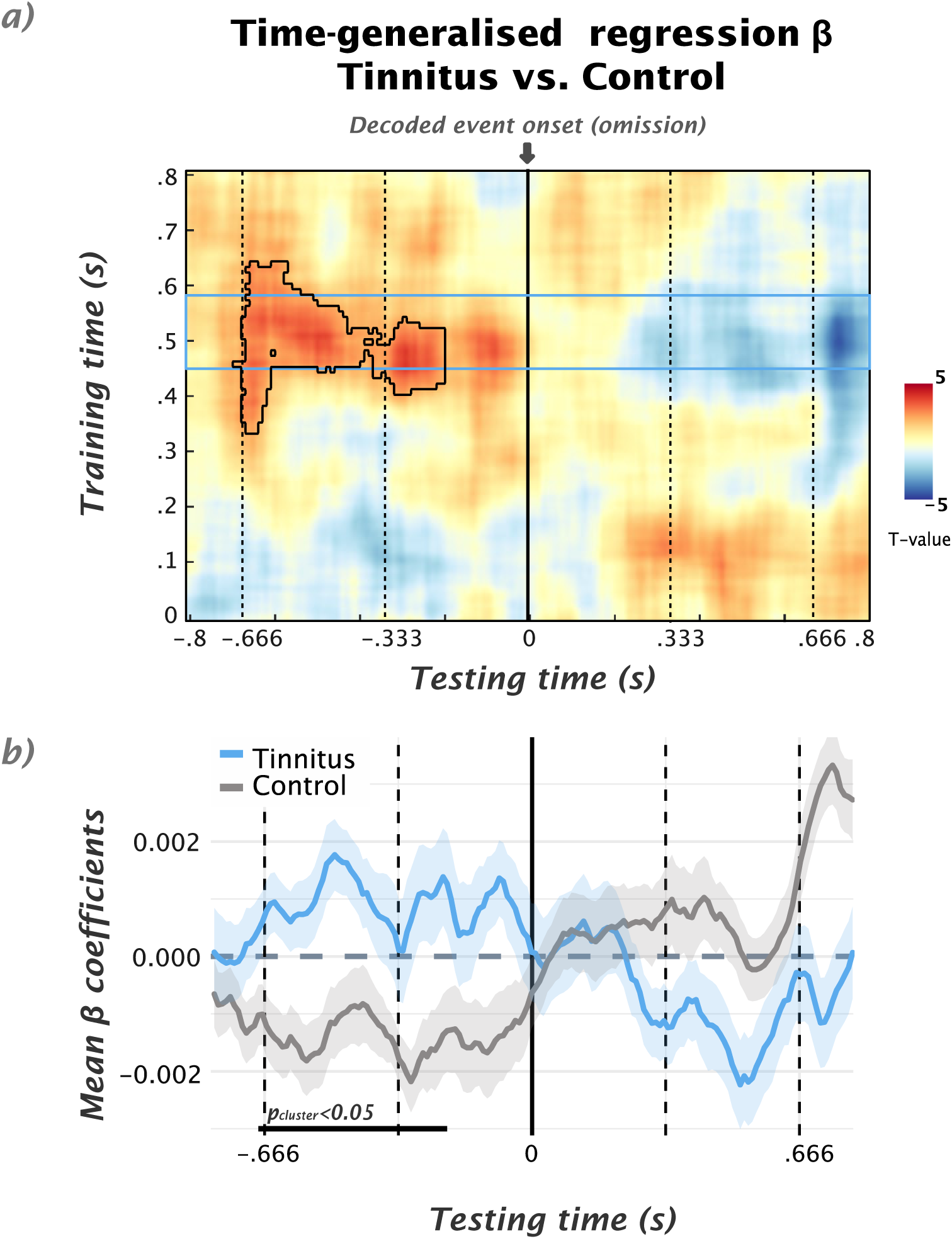
a) Group comparison (see Figure 1c) of β-coefficient values between Tinnitus vs. Control groups in time-generalised matrix in omission trials. Colors indicate t-values and solid black borders delimiting periods of significant difference (*p* < 0.05, cluster corrected). b) Time courses of β-coefficients averaged over 480-580ms training time-window (indicated by the blue rectangle and corresponding to the one previously demonstrated in sound-type trials), showing effects driven by a relative increase of regularity-dependent carrier frequency specific activity prior to anticipated onset period in Tinnitus group.

**Figure S5.**
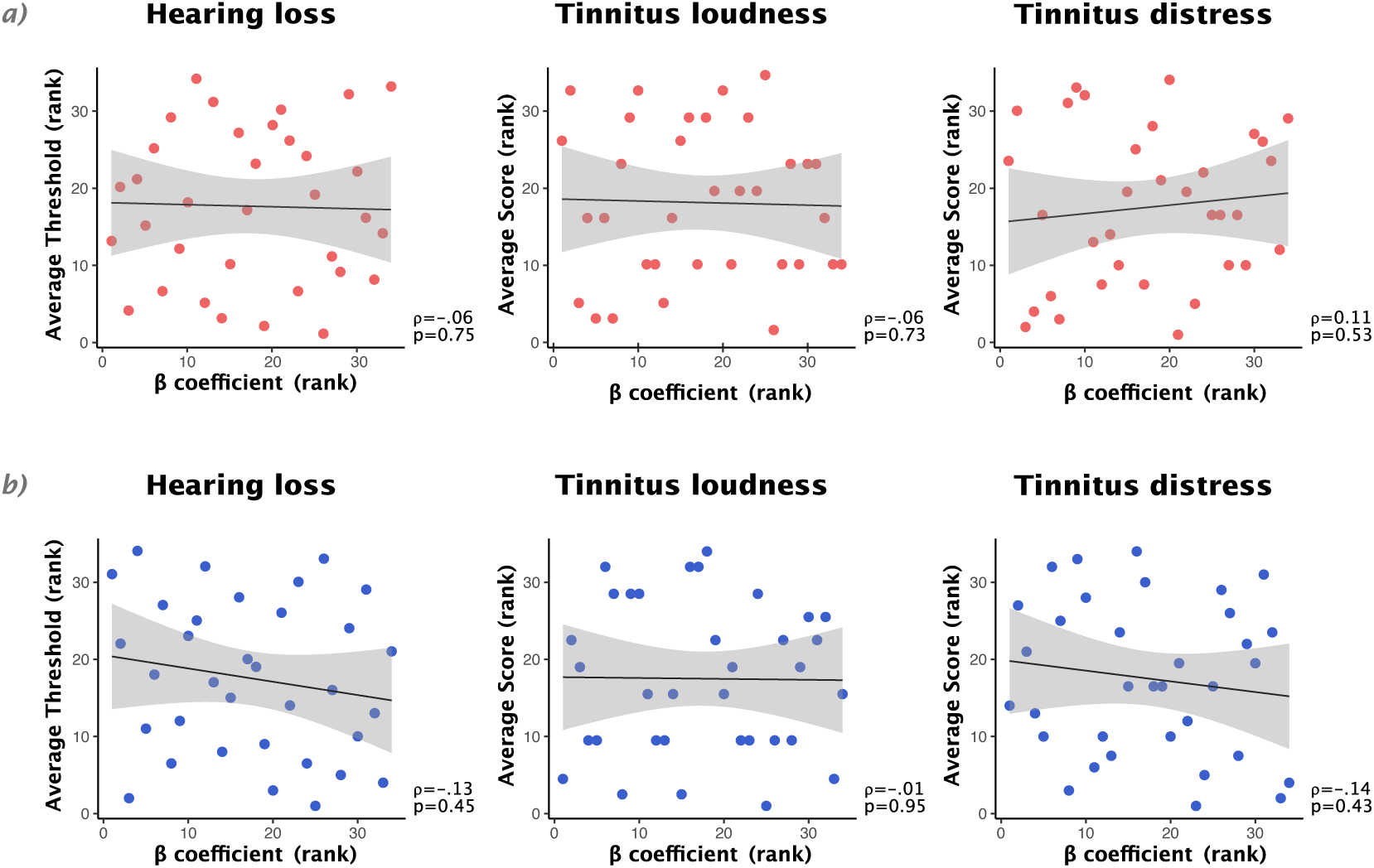
Scatter plots of hearing loss (left), tinnitus loudness (1-10 scale, middle) and tinnitus distress (TQ, right) measures with individual β-coefficient values (same as in Figure 2c) in Tinnitus group. a.) Pre-stimulus positive cluster, no significant correlation was revealed (*p* > 0.05, uncorrected). b) Post-stimulus negative cluster, no significant correlation was shown for any of the tested factors (*p* > 0.05, uncorrected).

